## Supplementary figures and images for "Investigating the native functions of [NiFe]-CODH through genomic context analysis"

### Supplementary File 5_Tree1.pdf

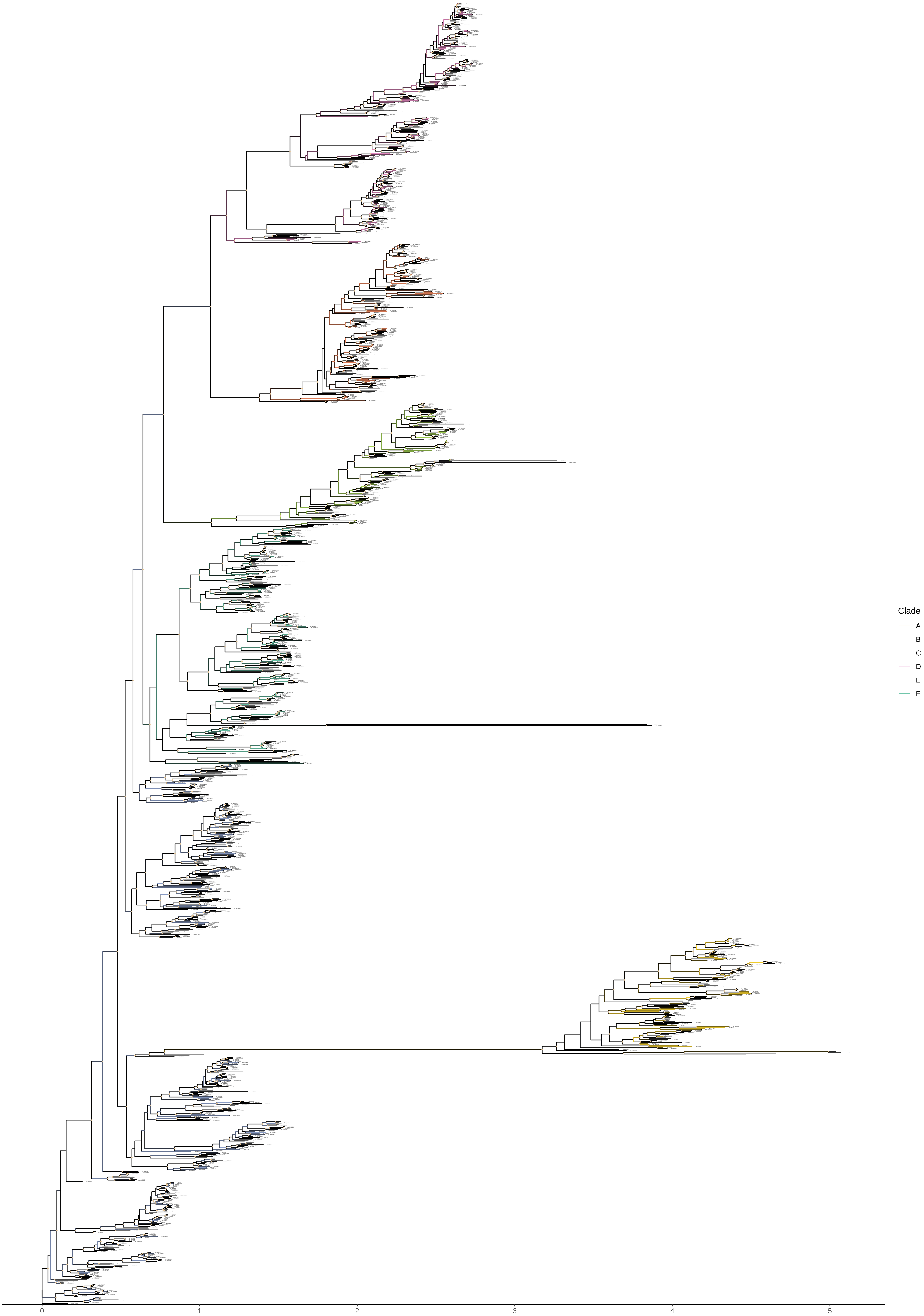

### Supplementary File 6_Tree2.pdf

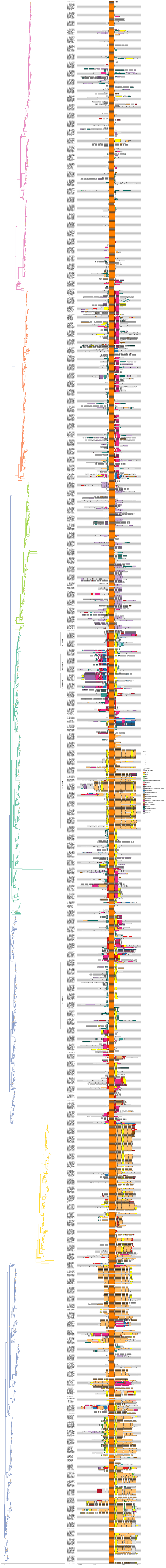

### Supplementary File 7_Tree3.pdf

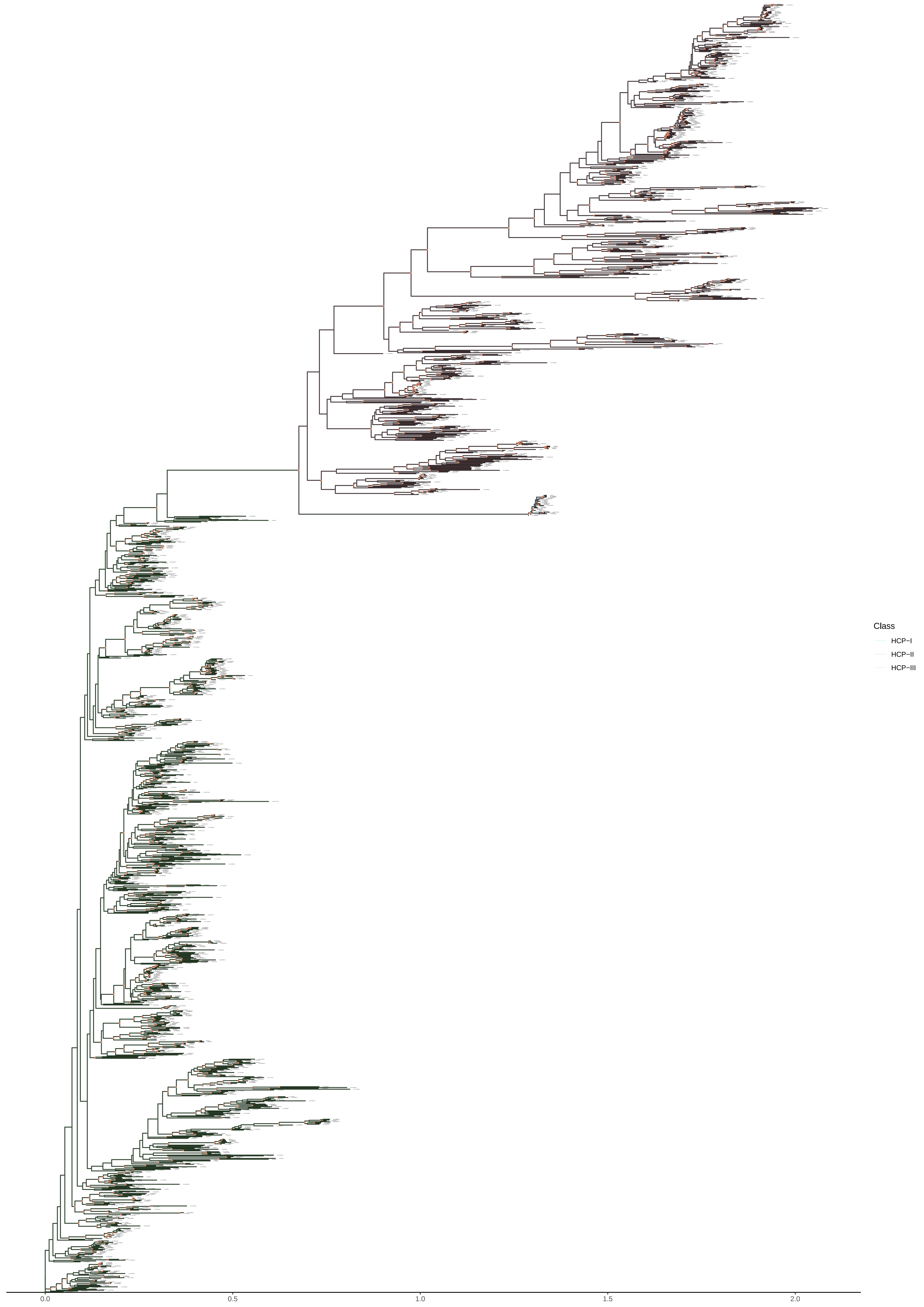

### Supplementary File 8_Tree4.pdf

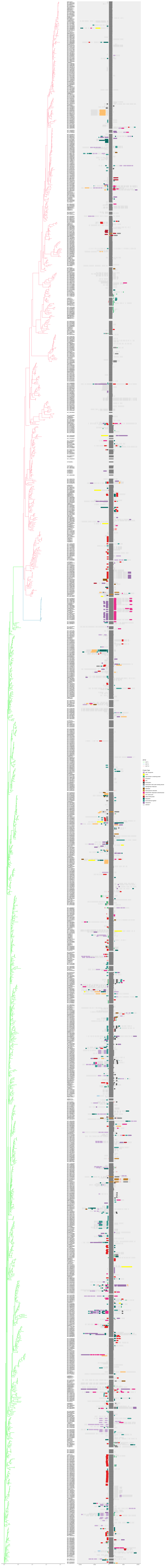

### Supplementary File 9_Tree5.pdf

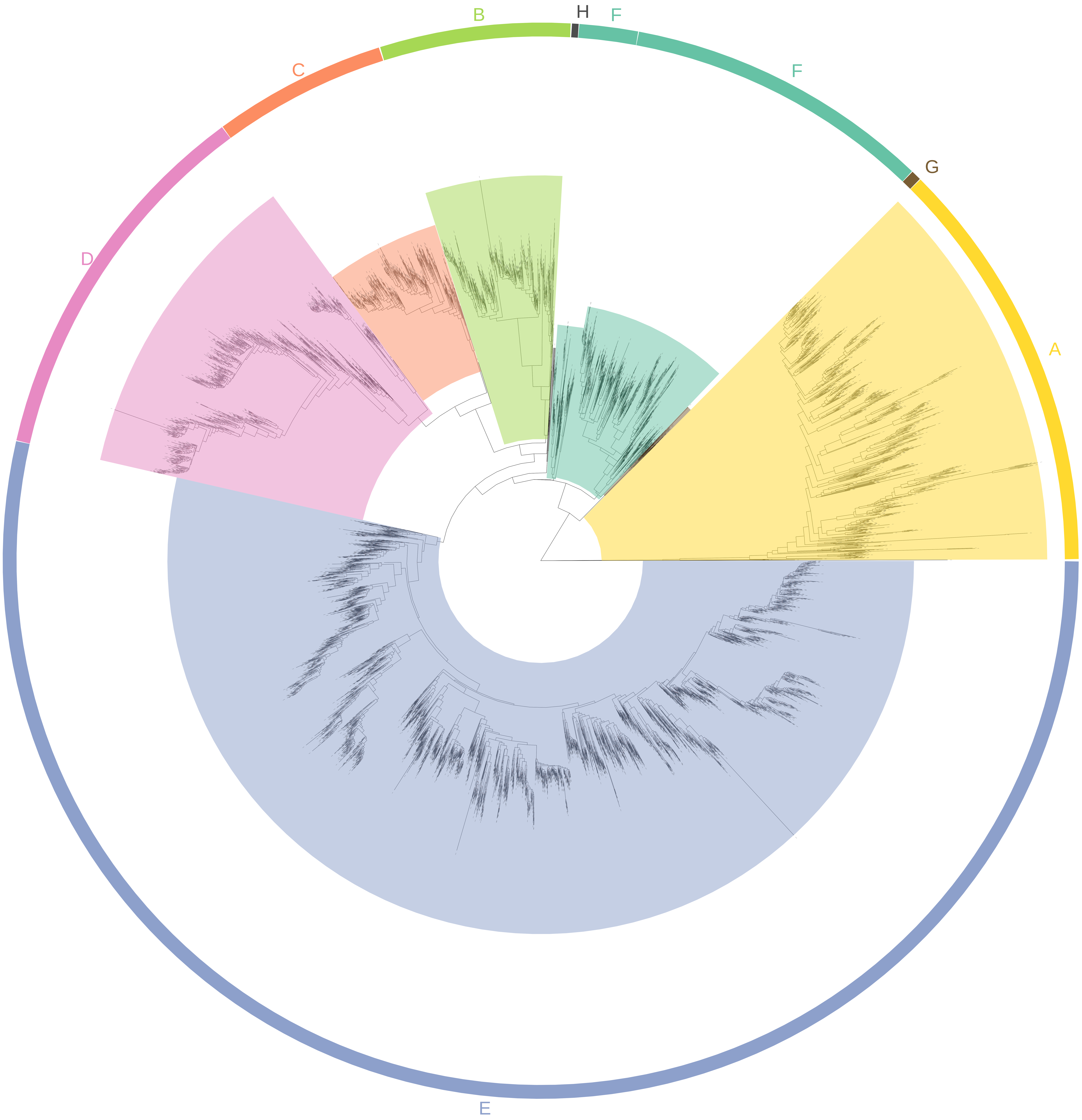

### Supplementary File 10_Tree6.pdf

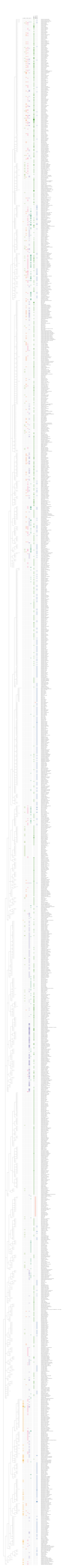

### Supplementary File 11_Tree7.pdf

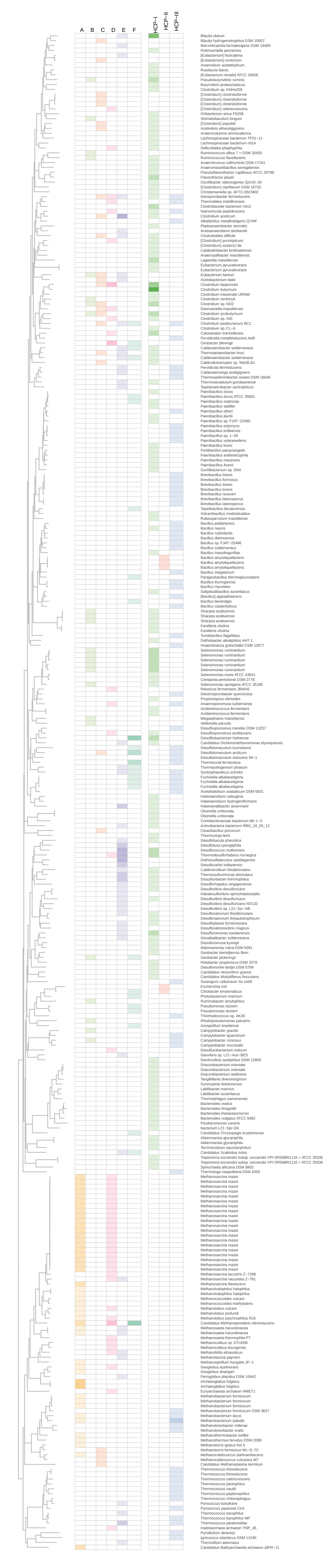
